# Inner visual experience biases object neural representational geometry during perceptual encoding

**DOI:** 10.1101/2025.08.01.668040

**Authors:** Shuang Tian, Xiaomin Mao, Dahui Wang, Xiaoying Wang, Yanchao Bi

## Abstract

Perception and mental imagery are widely thought to rely on overlapping neural representations, giving rise to perception-like visual experiences during imagery generation. However, such overlap, by itself, is agnostic to the nature of the underlying neural geometry for both perception and imagery. Here we addressed this question by comparing visualizers and individuals with congenital aphantasia using fMRI, representational similarity analysis, and multimodal deep neural network models to probe different types of neural geometry. Despite comparable behavioral performance and perception-imagery generalization of neural object representations in the ventral occipitotemporal cortex, the two groups exhibited systematically different object representational geometries already in the perceptual encoding phase. Visual-similarity-aligned geometry was selectively reduced in the left lateral occipitotemporal cortex of individuals with aphantasia during perceptual encoding, before imagery generation, and its expression during imagery covaried with imagery vividness. In contrast, language-relation-aligned geometry was preserved across groups during both perception and imagery. Together, these findings suggest that subjective imagery experience is associated with the geometry of neural object representations during both perception and imagery, and that representations within perception-imagery cross-decoding regions comprise heterogeneous components rather than a unitary shared neural code.

## Introduction

One can readily recognize a cat from visual input. One can also retrieve the appearance of a cat from memory (semantic knowledge) and mentally visualize it (“seeing with mind’s eye”) in the absence of sensory input (Fig. 1a). Decades of work have shown substantial overlap between the neural substrates of visual perception, visual semantic knowledge, and visual imagery, particularly in higher-order visual cortex, leading to the prevailing view that internally generated and externally driven visual representations rely on a common neural code (Barsalou et al., 2003; Martin, 2016; Pearson, 2019; Pearson & Kosslyn, 2015). Most studies have focused on demonstrating overlap between perception and imagery, asking whether the same neural populations are engaged in both processes (e.g., Harrison & Tong, 2009; Naselaris et al., 2015; Wadia et al., 2026). Much less is known about the representational content that supports this overlap. This question is difficult to address because perception, imagery, and object knowledge are normally tightly coupled.

**Fig. 1.**
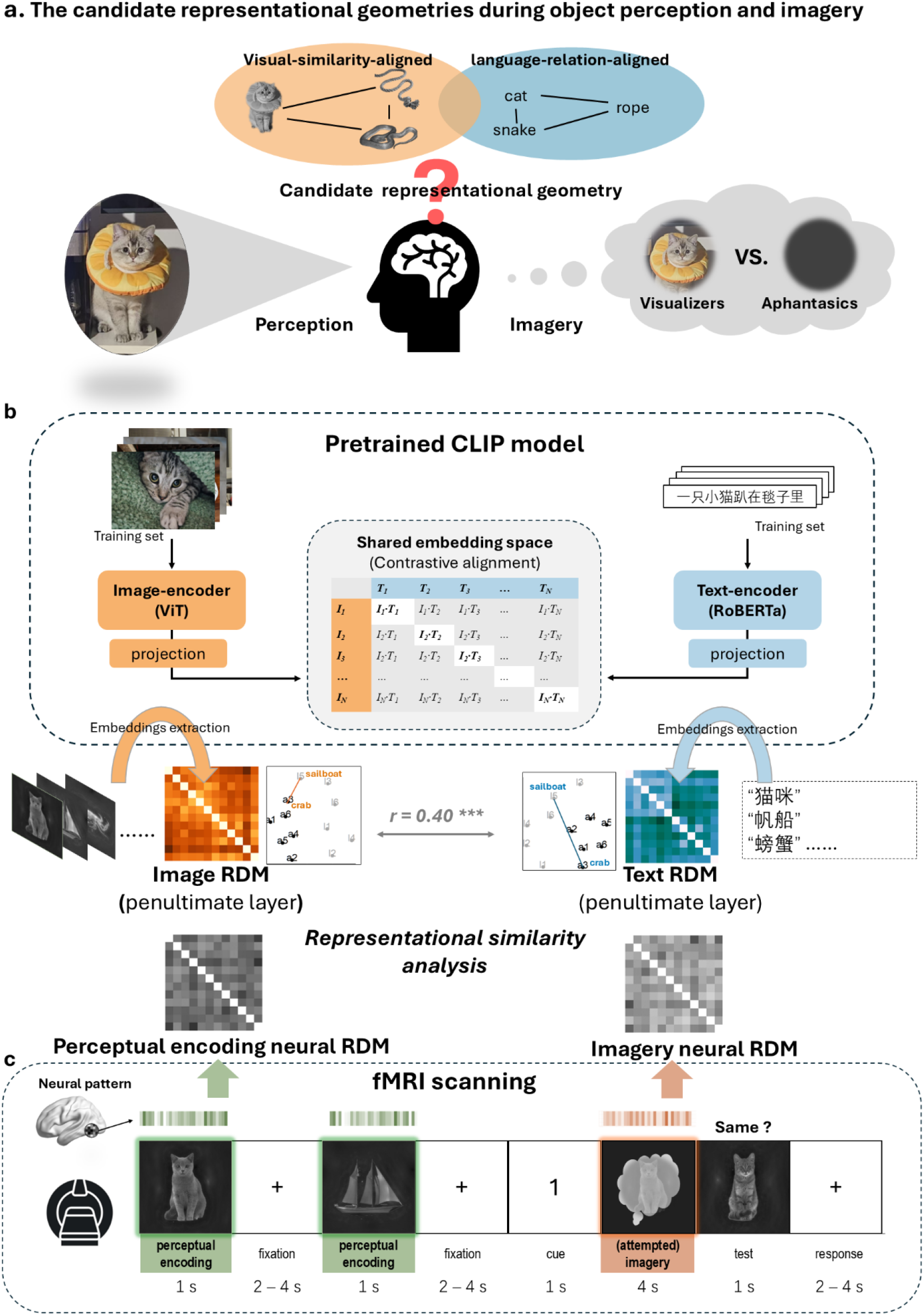
Study overview and analytical framework.a, We compared visualizers and individuals with aphantasia to examine whether and how imagery experience biases the neural representational geometry of objects during perception and imagery. **b,** Schematic illustration of the pretrained CLIP (Contrastive Language-Image Pretraining) model. The Chinese CLIP model was used to characterize candidate representational geometries. Example image-text pairs from the training set are shown for illustrative purposes only. Experimental images and their corresponding object labels were input to the image- and text- encoders, respectively. Embeddings from the penultimate layer (and all layers for the processing-stage analyses) were used to construct model representational dissimilarity matrices (RDMs) using cosine distance. The correlation between the penultimate-layer image- and text- encoder RDMs was r = .40 (p < .001). **c,** Retro-cue working memory paradigm used during fMRI scanning. Each trial began with sequential presentation of two images (perceptual encoding phase), followed by a numeric cue indicating which image to visualize (imagery phase). A test image was then presented, and participants judged whether it was exactly identical to the cued image. Neural activity patterns from the perceptual encoding and imagery phases were used to construct neural RDMs, which were subsequently compared with model- derived RDMs.

Congenital aphantasia provides a unique opportunity to dissociate them. These are individuals that report little or no conscious visual imagery, having “mind’s blindness” in a sense. The group shows largely preserved visual recognition, semantic knowledge, and many other forms of cognitive performance, with subtle differences in processing efficiency or cognitive strategy in some tasks (Keogh et al., 2021; J. Liu & Bartolomeo, 2023; Monzel, Vetterlein, et al., 2021; Zeman, 2024).This dissociation raises a fundamental question: what role does visual imagery experience play in shaping object representations in the human brain? If visual imagery is an integral component of how object knowledge is represented and maintained, then the absence of imagery experience may be associated with differences in the organization of neural object representations in broader processes, including, for example, perception. Conversely, if imagery is merely an epiphenomenal byproduct of object representations that are otherwise intact, then neural representational structure should remain largely unchanged.

Recent work on aphantasia has primarily examined neural representations during imagery in relation to perception. During attempted imagery generation Individuals with aphantasia manifest typical object domain-selective activity in the ventral visual cortices, similar to those observed in typical visualizers (J. Liu et al., 2025). However, in specific regions their neural activity pattern during attempted imagery is less similar to those during perception. For example, reduced correspondence of object representational similarity patterns between perception and imagery has been reported in the fusiform imagery node (FIN; J. Liu et al., 2025), and reduced decoding accuracy across imagery and perception for low-level visual features in early visual cortex (Chang et al., 2025). These findings were taken as evidence that imagery experience is supported by perception-like representations in these specific regions and not the other regions showing comparable imagery-perception decoding in the two groups. This interpretation implicitly assumes that perceptual representations provide a stable “visual” reference against which imagery representations are evaluated, and that the principal consequence of aphantasia lies in imagery-related processes rather than perception itself (Chang et al., 2025; Dijkstra et al., 2017; J. Liu et al., 2025).

However, emerging evidence suggests that perception itself may not be entirely invariant to aphantasia. Behavioral studies have reported differences in visual search, visual feature judgments, and object recognition (Liu & Bartolomeo, 2023; Monzel et al., 2021, 2023; Monzel & Reuter, 2024; see Zeman, 2024 for review). Neural differences during perception have also been observed, including altered activity strength in early visual cortex and altered connectivity between ventral visual regions and frontoparietal systems (Chang et al., 2025; J. Liu et al., 2025). These findings raise the possibility that subjective imagery experience is associated not only with imagery-specific processes, but also with the underlying representational organization of object knowledge expressed during perception. Under this view, successful behavioral performance and even shared decodability across perception and imagery may arise from different underlying representational structures.

Testing this possibility requires moving beyond demonstrations of shared activation or decodability to characterizing the content of neural representations with reference geometries. Research on aphantasia has explicitly proposed that individuals with aphantasia may rely more on verbal or semantic strategies in tasks that typically recruit visual imagery (Cabbai et al., 2023; Keogh et al., 2021; Monzel, Keidel, et al., 2021). This motivates language-derived semantic structure as a candidate organizing principle of object representations alongside visual similarity. Characterizing these candidate geometries is nontrivial because visual and language semantic structure are themselves strongly correlated, and because their relational spaces, particularly semantic ones, have historically been difficult to quantify. Recent advances in vision and language foundation models provide powerful tools for quantifying complex relational structures learned from visual and language experience (Chen et al., 2025; Shoham et al., 2024; Xu et al., 2025).

Multimodal vision-language models provide a particularly useful framework for studying visual object perception and subsequent imagery, where neural representations are initially triggered by visual inputs. Models such as CLIP (Fig. 1b; Radford et al., 2021; Yang et al., 2023) learn aligned representational spaces for images and language through large-scale contrastive learning. Unlike unimodal visual or language models, multimodal models explicitly learn correspondences between visual and language representations. As a result, they provide a framework in which object representations elicited by visual inputs can be evaluated with respect to both visual similarity-based and language-derived relational structures. Both vision and language models have exhibited representational similarity to neural representations in human visual cortices (Cichy et al., 2016; Doerig et al., 2025; Du et al., 2025), and language-supervised vision models further show unique contributions to explaining neural representations relative to unimodal vision models (Chen et al., 2025; A. Y. Wang et al., 2023). These properties make them particularly useful for testing whether neural representations associated with perception and imagery are better aligned with visual similarity or with the relational structures captured through language. If imagery experience contributes to the content of object representations, differences associated with aphantasia may emerge not only during imagery but also during the initial perceptual encoding of object information. Comparing neural representational geometry against these different model-derived reference spaces across perceptual encoding and imagery therefore provides a means of testing whether imagery experience is associated with systematic differences in the neural representation of object processing.

## Results

In this study, we took aphantasia as a natural imagery-experience-deprived population, examining the impact of imagery experience during both perceptual encoding and imagery maintaining of objects. We conducted an fMRI experiment using a retro-cue visual working memory paradigm (Fig. 1c) in individuals with and without visual imagery (visualizers vs. aphantasics).

We first identified brain regions with functional overlap (shared discriminability) across perceptual encoding and imagery phases by performing whole-brain searchlight SVM object domain decoding. This is a common approach for testing the shared code between perceptual and imagery-based neural representations (Chang et al., 2025; Harrison & Tong, 2009; Lee et al., 2012; Reddy et al., 2010), without revealing their specific representational contents.

Our key fMRI analyses were performed in these functionally overlapped regions, which have been the regions of interest (ROIs) in this study, to probe their representation contents. Visual-aligned vs. language-relation-aligned geometries derived from the multimodal deep neural network (i.e., CLIP model) were used as references to fit neural representations via linear mixed-effects RSA for the perceptual encoding phase and imagery phase separately. A follow-up searchlight RSA within the cross-phase-decoding regions was performed to explore which regions supported these representational geometries. Finally, we compared unimodal vision- and language- DNN models with the multimodal CLIP encoders across multiple layers in representational similarity to the cross-phase-decoding regions, to examine the importance of multimodal alignment.

Note that whole-brain searchlight RSA using the CLIP encoders revealed no significant clusters showing imagery experience effects surviving correction for multiple comparisons outside these predefined ROIs.

## Behavioral results: Comparable behavioral performance in visualizers and aphantasics

To test whether aphantasics could perform the retro-cue working memory task as well as visualizers, we analyzed participants’ behavioral performance in the scanner. As shown in Fig. 2a, a repeated-measures ANOVA on accuracy revealed no significant main effect of group (F_(1, 37)_ = 1.36, p = .25, η²_p_ = .04), stimulus domain (F_(1, 37)_ = 0.19, p = .38, η²_p_ = .005), or their interaction (F_(1, 37)_ = 0.67, p = .42, η²_p_ = .02).

**Fig. 2.**
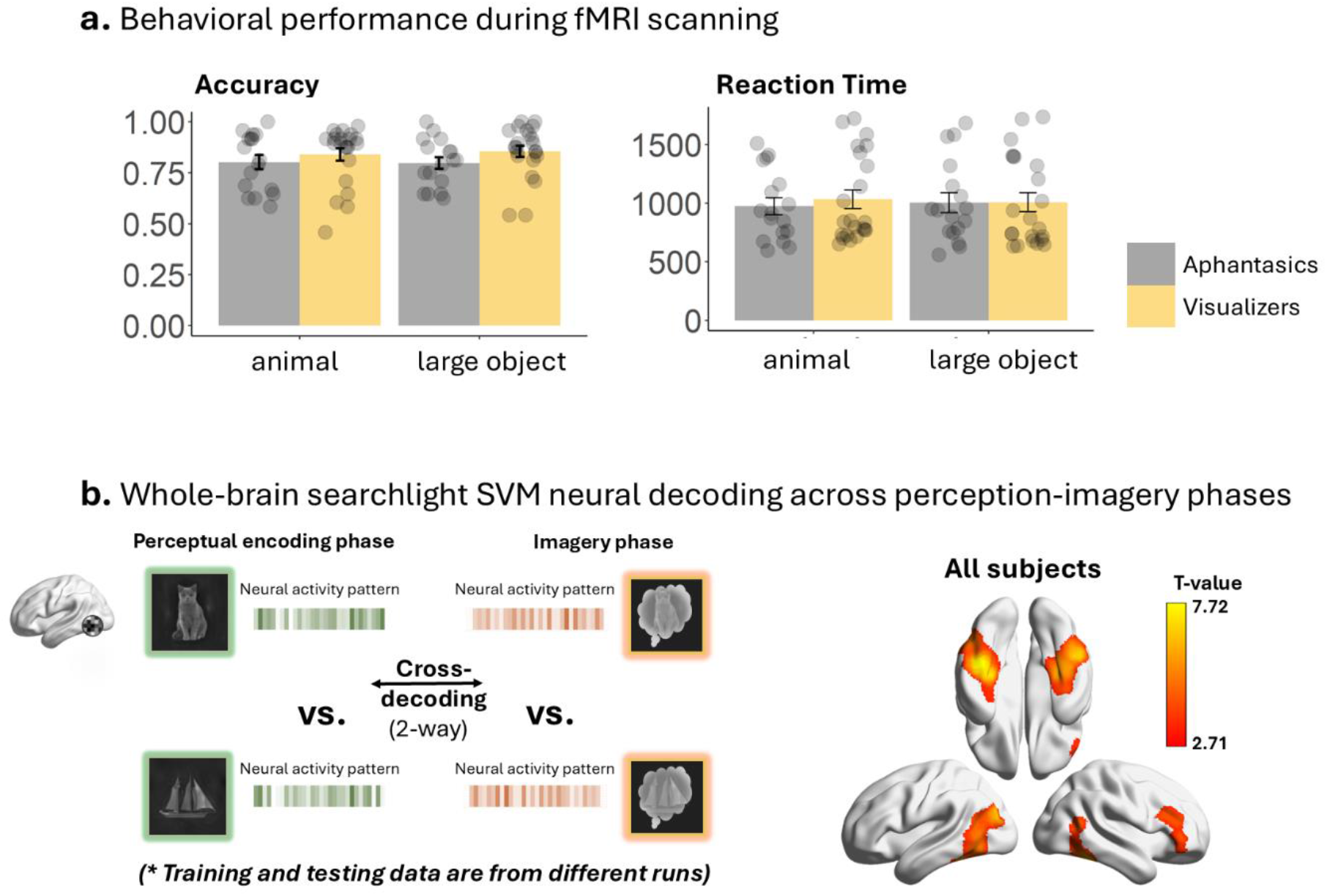
Behavioral performance and cross-phase decoding results. a, Behavioral performance. Bars show mean accuracy and reaction times (RTs) for visualizers (yellow) and individuals with aphantasia (gray). Error bars indicate ±1 SEM, and dots represent individual participants. **b,** Searchlight SVM cross-phase decoding results. Searchlight SVM domain-level (animal vs. large object) cross-phase-decoding map across all participants (voxelwise p < .001, cluster-level FWE-corrected p < .05). No significant group differences were observed; uncorrected group comparison maps are shown in Supplementary Fig. 2. This map defined regions showing shared perception-imagery representations and served as the regions of interest for subsequent representational similarity analyses. Item-level cross-phase decoding results are provided in Supplementary Fig. 3.

For reaction time (RT), a repeated-measures ANOVA showed no significant main effect of group (F_(1, 37)_ = 0.08, p = .78, η²_p_ = .002) or stimulus domain (F_(1, 37)_ = 0.02, p = .88, η²_p_ = .0006). Although the interaction was significant (F_(1, 37)_ = 5.43, p = .03, η²_p_ = .13), post-hoc simple effects were not significant (ps > .08).

These results indicate that aphantasic participants performed the task as accurately and efficiently as visualizers.

## Localizing the regions of interest by successful SVM decoding across the perceptual encoding and imagery phases

We focused on brain regions supporting (at least some) functional overlaps between perceptual encoding and imagery phases, by conducting whole-brain searchlight SVM cross-phase-decoding analyses for object domains (i.e., animal vs. large object 2-way classification). A repeated split-half cross-validation approach was used, where an SVM classifier was trained on data of perceptual encoding from half of the runs and tested on imagery data from the remaining half, and vice versa. For each searchlight sphere, the decoding accuracy was averaged across all split-half combinations and both train-test directions, and the resulting value was assigned to the center voxel. This procedure was repeated across a grey matter mask (see Methods). Group-level statistical inference was conducted using permutation test. Results were thresholded at voxelwise p < .001, cluster-level FWE-corrected p < .05, unless stated otherwise.

As shown in Fig. 2b, group-level analysis combining all subjects yielded significant cross-phase decoding in the bilateral ventral occipitotemporal cortex (VOTC), bilateral posterior middle temporal gyrus (pMTG), with extensions into the left lateral occipital cortex (LOC), as well as in the right inferior frontal gyrus (IFG). The group-level result of each group (Supplementary Fig. 2) showed that the visualizer group had significantly successful cross-decoding for domains in bilateral VOTC, extending to the left LOC. The aphantasic group showed significant cross-decoding in the right fusiform gyrus (FG). All of these areas located in the regions identified by the combined group-level result. We then tested group differences within these regions. The result showed a trend toward higher accuracy in the left ventral lateral occipitotemporal cortex (LOTC) in visualizers, consistent with previous literature (J. Liu et al., 2025; Spagna et al., 2021), although this effect did not survive multiple comparisons correction (see Supplementary Fig. 2 for an uncorrected result).

Item-level cross-phase decoding (12-way classification) for real objects revealed similar but spatially more restricted results (Supplementary Fig. 3) than domain-level cross-phase decoding. Group-level result of all subjects showed a distribution across VOTC that largely overlapped with the domain-level decoding result. The visualizer group showed significant cross-decoding in the left FG, and the aphantasic group showed significant cross-decoding in the right lateral FG. Again, no significant group differences were observed after correction (see Supplementary Fig. 3 for an uncorrected result).

These results suggest that the VOTC regions contained shared object-discriminative representations across the perceptual encoding and imagery phases in both groups, with the right IFG additionally showing shared domain-discriminative representations. Critically, successful cross-phase decoding alone does not reveal underlying representational structures, nor whether they are the same across different groups. Therefore, we performed the following RSA to disentangle potentially different types of neural representational geometry in these regions (Fig. 2b), using both an ROI-based approach and a small-volume correction approach (i.e., searchlight within the regions).

## Representational geometry during the perceptual encoding phase

### Visual- vs. Language-relation- biased representational geometry construction

To probe the contents of the neural representations, we leveraged a pre-trained Chinese version of the multimodal deep neural network CLIP (Yang et al., 2023), which consists of a text-encoder and an image-encoder that were trained to align their embedding spaces. We input the stimuli images into the image-encoder and the names of the objects into the text-encoder, extracted the embeddings from the *penultimate layers*, which preserve modality-biased structures and maintain objective-relevant high-level representations (see results of earlier layers in the Models Comparison section below). These embeddings were computed and transformed into RDMs based on cosine distance (Fig. 1b). As expected, the CLIP image-encoder RDM and the text-encoder RDM had medium correlation (Pearson’s r = 0.40, p < .001), and the following analyses dissociate their specific effects statistically using linear mixed-effects model and partial correlation analyses.

### ROI-based RSA results

Using all regions showing significant cross-phase decoding (Fig. 2b) as a single ROI, we conducted RSA to examine the underlying representational geometry of the neural RDM and whether it is affected by visual imagery experience during the perceptual encoding phase. To estimate each encoder’s unique contribution and its interaction with group membership, as well as handle subject-level variance, we conducted linear mixed-effects analysis for this ROI-based test. A linear mixed-effects model was constructed to predict neural RDMs from two hypothesized representational geometries (CLIP image- and text-encoder RDMs derived from the penultimate-layer embeddings) and subject group, with subject included as a random effect. The full model was specified as: Neural RDM ∼ 1 + (CLIP image-encoder + CLIP text-encoder) * Group + (1 + CLIP image-encoder + CLIP text-encoder | subject). The model was simplified when necessary to address convergence issues (see Methods for details). The model allows for uncovering three main effects (CLIP-image encoder, CLIP text-encoding, group) and two interactions (CLIP-image encoder × group, CLIP-text encoder × group) in predicting neural RDMs.

During the perceptual encoding phase, there were significant main effects of the CLIP text-encoder (β = 0.67, p < .001) and group (β = −0.06, p < .001), with no significant interaction between these two. This suggests that although representational dissimilarity values were overall lower in visualizers, CLIP text-based alignment did not differ between the two groups. For image encoder, its main effect was not significant but its interaction with group was (β = 0.22, p = .023; see Table 1 for full statistics). Post hoc simple slope analyses confirmed that the CLIP image-encoder effect was significantly stronger in visualizers than in aphantasics (see Fig. 3a for the visualization of simple slopes).

**Fig. 3.**
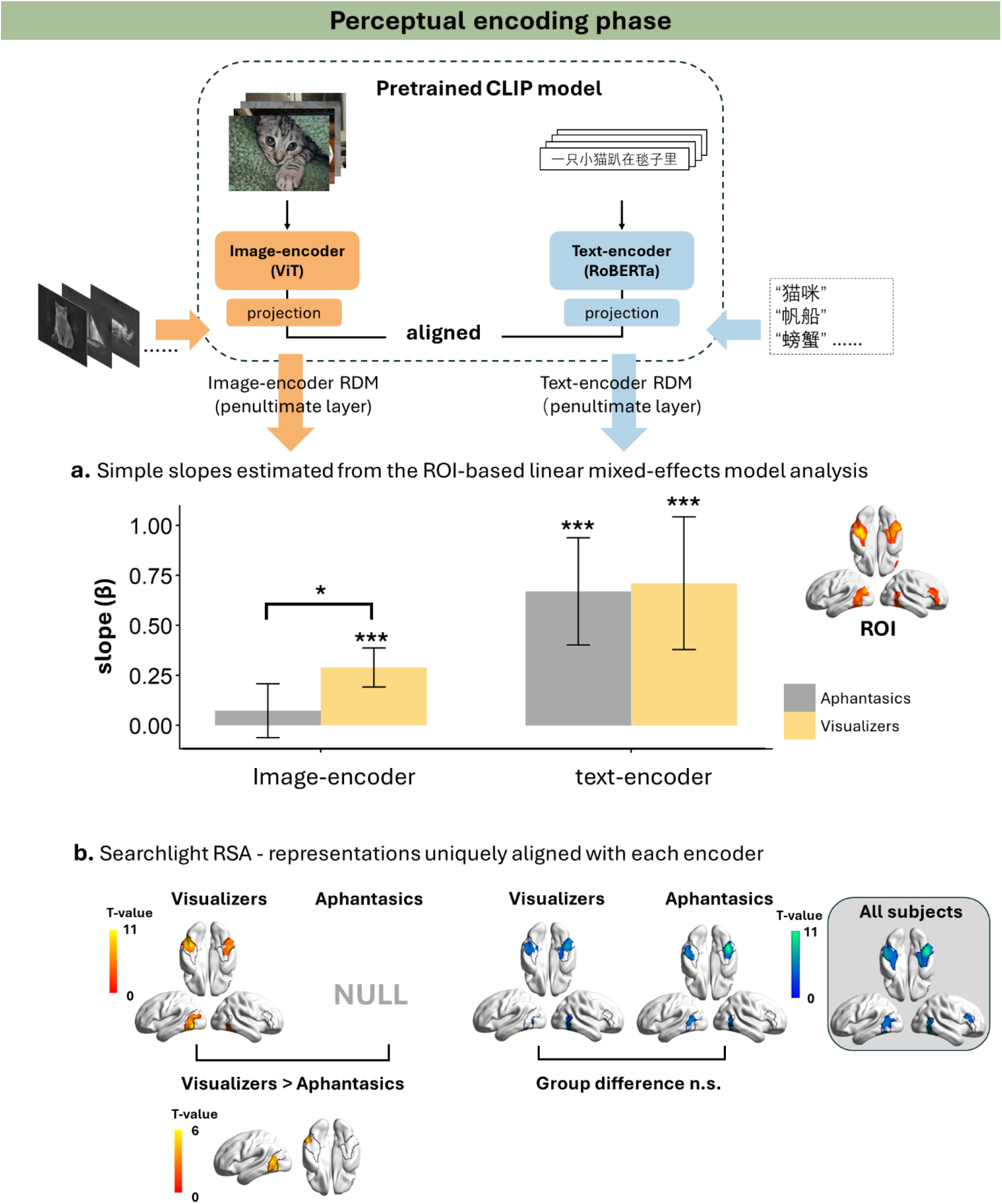
Representational geometry during the perceptual encoding phase. Left column: RSA with the CLIP image-encoder. **Right column:** RSA with the CLIP text-encoder. **a,** ROI-based linear mixed-effects model analysis. The ROI was defined from the searchlight SVM cross- phase-decoding map (Fig. 2b). Bars represent simple slopes (β) for each group estimated from the linear mixed-effects model. Error bars indicate 95% CI. Significance markers: #p<.1; *p < .05; **p < .01; ***p < .001. **b,** Searchlight RSA within the cross-phase-decoding mask (outlined in black; Fig. 2b). For each voxel (i.e., the center of each searchlight sphere), partial correlations were computed between the neural RDM and one CLIP encoder RDM while controlling for the other encoder RDM. When no significant group differences were observed, group-level analyses of all subjects (i.e., combining the visualizer and aphantasic groups) were performed to enhance sensitivity to shared effects; results from these analyses are shown in the gray boxes. Statistical maps were thresholded at voxelwise p < .001 and cluster-level FWE-corrected p < .05.

**Table 1.**
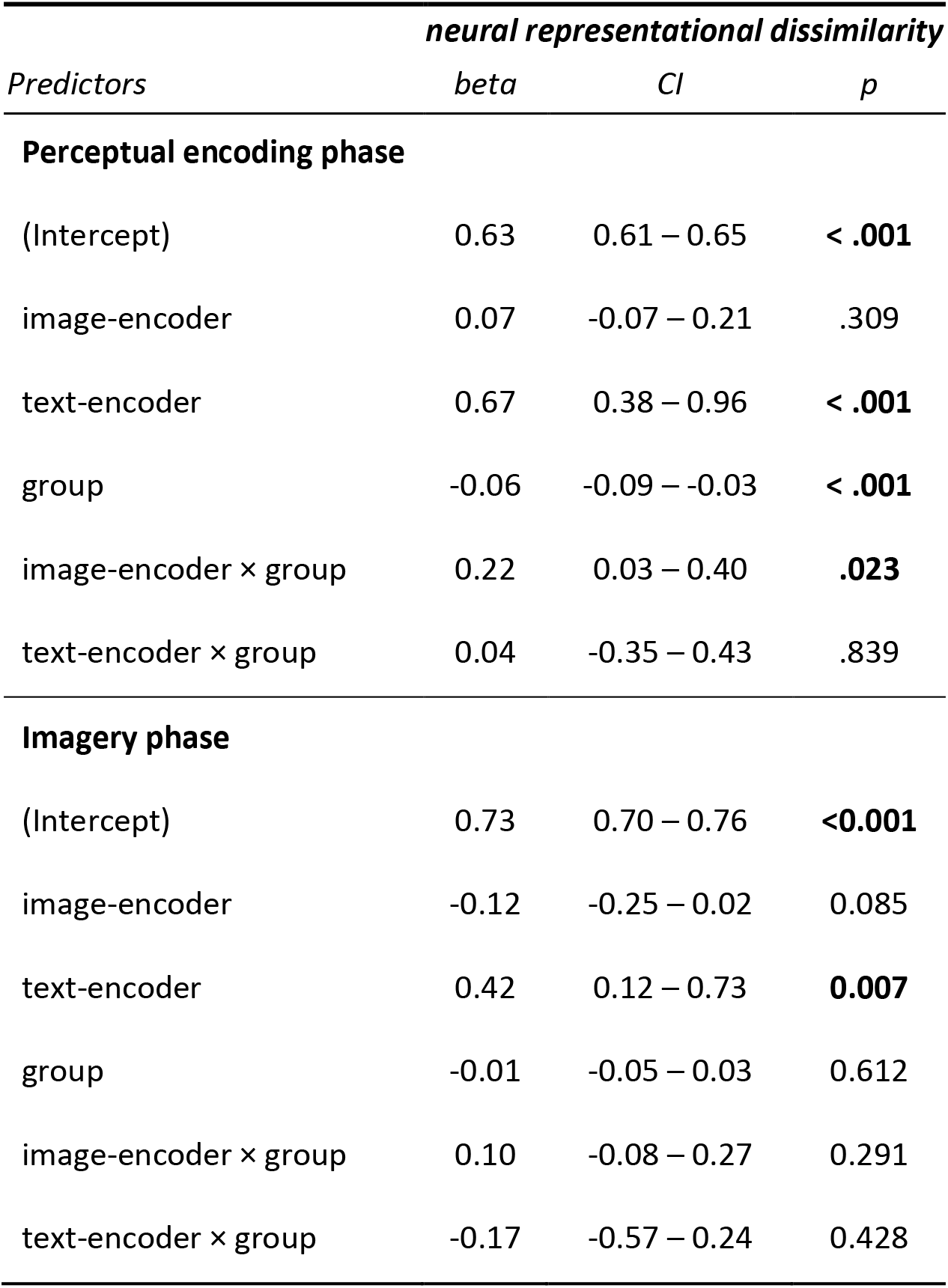
Results of linear mixed-effects model analysis of RSA

That is, both groups exhibited comparable text-encoder-specific representational geometry, whereas specific similarity to the image-encoder geometry was modulated by imagery experience, with individuals with aphantasia showing reduced similarity.

### Searchlight RSA within the cross-phase-decoding regions

To examine whether representational geometry differed across brain regions, we performed a searchlight RSA within the ROI (Fig. 2b). To ensure computational efficiency and avoid the frequent model convergence failures associated with fitting complex linear mixed-effects models across thousands of individual voxels, we simplified the small-volume searchlight-based analysis by direct correlation analyses. Given our interest in the unique contributions of each CLIP encoder to neural representations, the main results describe the partial correlations. Results were thresholded at voxel-wise p < .001, cluster-level FWE-corrected p < .05, unless stated otherwise. Uncorrected results are shown in Supplementary Figs. 4-5. Results of zero-order correlation of each encoder and commonality analysis (i.e., the shared variance among the neural, image-encoder, and text-encoder RDMs) are shown in Supplementary Fig. 6.

In visualizers, searchlight RSA revealed unique effects of the CLIP image-encoder (penultimate layer; Fig. 3b, left) within bilateral VOTC, extending into the left LOC. No significant unique effects of the image-encoder were detected in aphantasics. A direct group comparison test revealed a significant difference in the left LOTC.

By contrast, searchlight RSA with the CLIP text-encoder (penultimate layer; Fig. 3b, right) showed that both groups had significant partial correlations in bilateral FG and pMTG, without regions showing significant group differences. Thus, we combined all subjects to increase power for detecting the CLIP text-encoder-specific effect, yielding significant effects in bilateral FG, bilateral pMTG, and right IFG (statistical maps in the grey box).

These findings corroborate and extend the ROI-based linear mixed-effects model results. Across both analyses, visualizers exhibited stronger neural representational alignment with the CLIP image-encoder geometry than individuals with aphantasia, indicating an imagery-experience effect. The searchlight analysis further localized this group difference to the left LOTC. In contrast, both analyses consistently revealed significant and comparable unique alignment with the CLIP text-encoder geometry in the two groups, with the searchlight analysis localizing these effects to the bilateral FG, bilateral pMTG, and right IFG.

## Representational geometry during the imagery phase

The imagery phase analyses mirrored the pipeline used in the perceptual encoding phase: 1) ROI-based linear mixed-effects model RSA to examine the effects of CLIP encoders and their interactions with subject groups; 2) a small-volume searchlight RSA to identify the specific brain regions showing effects of CLIP encoders.

### ROI-based RSA results

During the imagery phase (Fig. 4a), the linear mixed-effects model RSA of the cross-phase-decoding ROI showed that only the main effect of the CLIP text-encoder was significant (β = 0.42, p = .007), with no significant interaction with group (β = -0.17, p = .43; see Fig. 4a for simple slopes). Neither the main effects of the CLIP image-encoder or group, nor their interaction, reached significance (all ps > .08; see Table 1 for full statistics). These results indicate that neural RDMs in both groups exhibited comparable unique alignment with the CLIP text-encoder geometry during imagery.

**Fig. 4.**
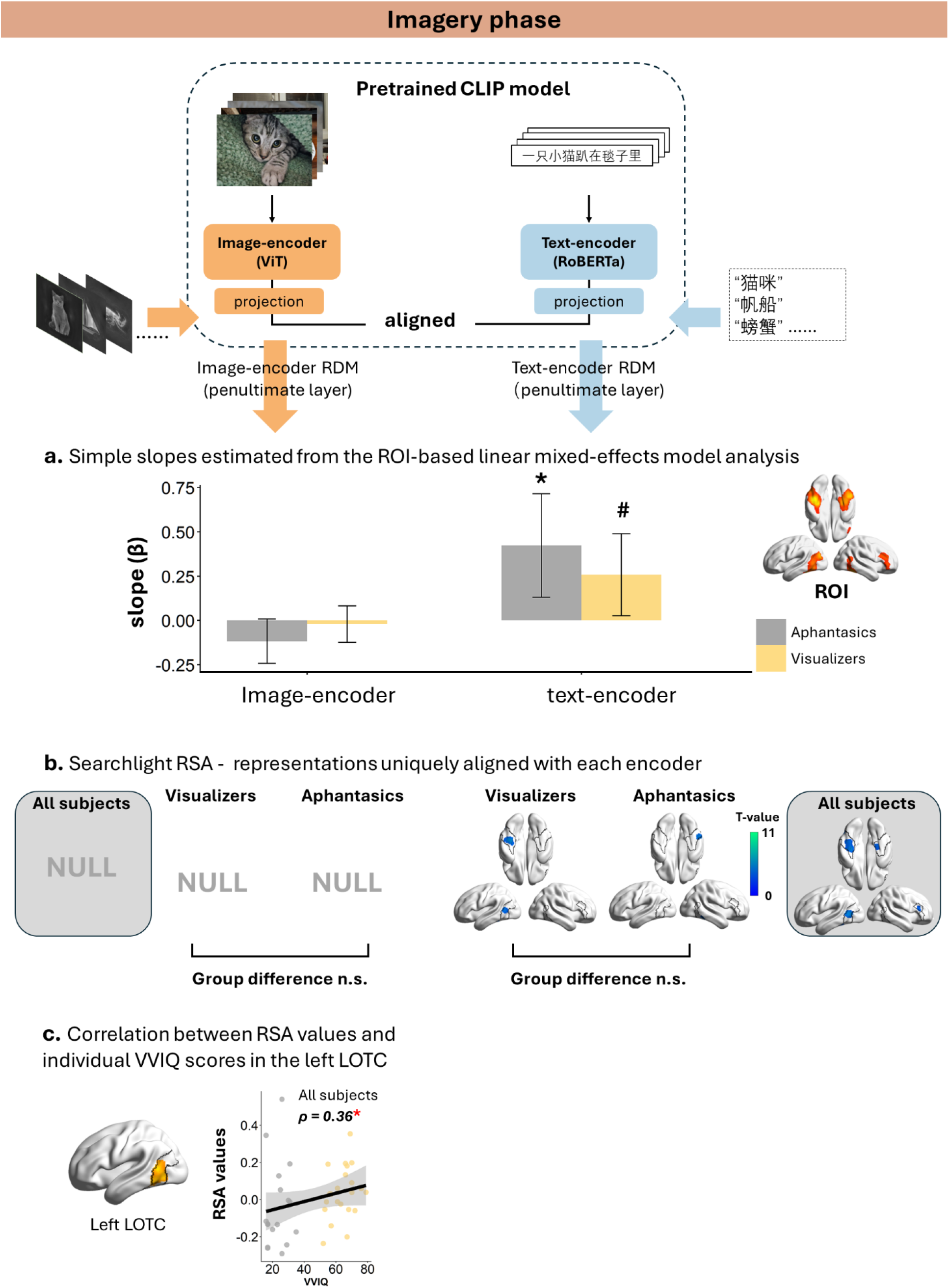
Representational geometry during the imagery phase. Left column: RSA with the CLIP image-encoder. **Right column:** RSA with the CLIP text-encoder. **a,** ROI-based linear mixed-effects model analysis. The ROI was defined from the searchlight SVM cross- phase-decoding map (Fig. 2b). Bars represent simple slopes (β) for each group estimated from the linear mixed-effects model. Error bars indicate 95% CI. Significance markers: #p<.1; *p < .05; **p < .01; ***p < .001. **b,** Searchlight RSA within the cross-phase-decoding mask (outlined in black; Fig. 2b). For each voxel (i.e., the center of each searchlight sphere), partial correlations were computed between the neural RDM and one CLIP encoder RDM while controlling for the other encoder RDM. When no significant group differences were observed, group-level analyses of all subjects (i.e., combining the visualizer and aphantasic groups) were performed to enhance sensitivity to shared effects; results from these analyses are shown in the gray boxes. Statistical maps were thresholded at voxelwise p < .001 and cluster-level FWE-corrected p < .05. **c**, Correlation between RSA values of the left LOTC and VVIQ scores across subjects. The ROI of the left lateral occipitotemporal cortex (LOTC) was defined from the searchlight RSA group difference map during the perceptual encoding phase (Fig. 3b, left-bottom). RSA values in the left LOTC were the partial correlations between neural RDMs during the imagery phase and the CLIP image-encoder penultimate- layer RDM while controlling for the text-encoder RDM.

The separate analyses of the perceptual encoding and imagery phases revealed different patterns of neural representational geometry. To formally test whether the encoder × group interaction differed between phases, we fitted an additional linear mixed-effects model including task phase as a factor, thereby testing the three-way interaction among encoder, group, and task phase. The three-way interaction was not significant for either the CLIP image-encoder (β = 0.12, p = .22) or the CLIP text-encoder (β = 0.21, p = .34). Thus, although the encoder × group interaction reached significance only during the perceptual encoding phase for the CLIP image-encoder, the absence of a significant three-way interaction indicates that differences between the two task phases should be interpreted with caution.

### Searchlight within the cross-phase-decoding regions

During the imagery phase, searchlight RSA within the cross-phase-decoding regions with the CLIP image-encoder revealed no significant clusters in either group, nor any group differences; no significant clusters were observed when all subjects were combined to boost potentially group-consistent effects, either. For the CLIP text-encoder (Fig. 4b, right), the visualizer group showed significant unique effects in the left FG and left pMTG; the aphantasic group showed the unique effects in the right lateral FG. Comparing the two groups revealed no significant clusters at the corrected threshold. The group-combined results (Fig. 4b; statistical maps in the grey box) showed significant CLIP text-encoder-unique effects in bilateral FG, left pMTG, and right IFG, largely overlapping with those observed during the perceptual encoding phase.

### Correlation between individual vividness and visual-aligned geometry in the left LOTC

The left LOTC in the perceptual encoding phase manifested interesting subject group differences: Those without imagery experience respected less of the CLIP image-encoder-unique geometry. Such group differences were not significant in the imagery phase. Given the possibility that this null result in the imagery phase was due to substantial individual variability in subjective imagery vividness, we further examined the effect of imagery experience by leveraging continuous variations in imagery vividness. We took the left LOTC as the ROI (i.e., shown in Fig. 3b left bottom), examined the correlation between its RSA values and the continuous VVIQ scores (Vividness of Visual Imagery Questionnaire; Marks, 1973) across subjects during the imagery phase. RSA values were computed as the partial correlation between neural RDMs and the CLIP image-encoder RDM constructed from the penultimate-layer embeddings, controlling for the CLIP text-encoder RDM, as in the preceding analyses.

During the imagery phase, VVIQ scores were positively correlated with RSA values reflecting the unique alignment between neural RDMs and the CLIP image-encoder geometry in the left LOTC (Spearman’s ρ = 0.36, p = .02; Fig. 4c) across all subjects. The correlation within each group separately did not reach significance likely due to reduced sample size (visualizer group: Spearman’s ρ = 0.32, p = .15; aphantasia group: Spearman’s ρ = 0. 071, p = .76, uncorrected).

Together, the results indicate that how neural representations respected the CLIP image-encoder geometry is indeed affected by imagery experience, showing categorical group differences during the perceptual encoding phase, and scaling with subjective imagery vividness during the imagery phase.

## Comparing CLIP encoders and unimodal DNN models

Previous studies have shown that multimodal DNN models provide unique explanatory power for neural representations beyond that provided by unimodal models during visual perception (Chen et al., 2025; A. Y. Wang et al., 2023). To assess whether the contribution of multimodal alignment extends across different task phases and participant groups, we further compared multimodal and unimodal DNN models in deriving representational geometries that explains the neural representations of the cross-phase-decoding ROI (Fig. 2b). We considered models that share the same transformer backbones as the CLIP encoders but were trained exclusively on images or texts, i.e., unimodal vision and language models (DINOv2 for vision and RoBERTa for text; Y. Liu et al., 2019; Oquab et al., 2024). Because high-level representations may emerge at different processing stages across DNN models due to their distinct training objectives (Kornblith et al., 2021; Yamins & DiCarlo, 2016; Yosinski et al., 2014), we performed comparisons across multiple layer-groups (early, middle, late, divided evenly in each model). As this analysis aimed to test the functional consequences of multimodal alignment, we examined the ROI-based RSA results of each model without partialing out the alternative models, and carried out three-way repeated-measures ANOVAs with model type (i.e., unimodal vs. multimodal), layer-group, and subject-group on the Fisher’s Z-transformed correlation coefficients.

During the perceptual encoding phase, for vision models, the three-way repeated-measures ANOVA revealed a significant three-way interaction (F_(2,74)_ = 4.00, p = .04, η^2^ = 0.10; see Supplementary Table 1 for full statistics). Simple-effects analyses showed a significant main effect of model only in the middle layer-group (F_(1,37)_ = 19.50, p = 7.67 × 10^-4^, FDR-corrected), with DINOv2 exhibiting higher values than the CLIP image-encoder. For language models, the three-way ANOVA revealed a significant interaction between model type (i.e., CLIP text-encoder vs. RoBERTa) and layer-group (F_(2,74)_ = 39.20, p = 2.46 × 10^-12^, η^2^ = 0.51). Simple-effects analyses showed that RoBERTa yielded higher correlations than the CLIP text-encoder in the early and middle layer-groups (Fs > 9.19, ps < .004, FDR-corrected), whereas this pattern reversed in the late layer-group (F_(1,38)_ = 19.20, p = 1.33 × 10^-4^, FDR-corrected).

During the imagery phase, for vision models, the three-way ANOVA revealed only a main effect of model (F_(1,37)_ = 4.66, p = .04, η^2^ = 0.11), with the CLIP image-encoder showing higher RSA values than DINOv2. For language models, no significant effects but a trend toward a model difference in the late layer-group (CLIP text-encoder > RoBERTa, Welch’s t_(75.3)_ = 2.16, p = .03, uncorrected) were observed (see Supplementary Table 1).

Together, these results indicate that unimodal models accounted for neural representations during the perceptual encoding phase and even outperformed their multimodal CLIP counterparts at specific processing stages. By contrast, during the imagery phase, the unimodal models consistently showed weaker correspondence with neural representations than the multimodal CLIP models, suggesting that multimodal representational alignment becomes important when representations must be internally reconstructed in the absence of direct sensory input.

## Discussion

The present study investigated whether individual differences in visual imagery experience are reflected in the neural representational organization of objects during both perception and imagery. Previous work has demonstrated substantial overlap between perception and imagery, which has often been interpreted as evidence that the two processes rely on a shared neural code (Dijkstra et al., 2017; Harrison & Tong, 2009; Lee et al., 2012; Reddy et al., 2010; Wadia et al., 2026). Using model-based representational analyses within perception-imagery cross-decoding regions, we showed a more nuanced picture, as summarized in Fig. 6.

**Fig. 5.**
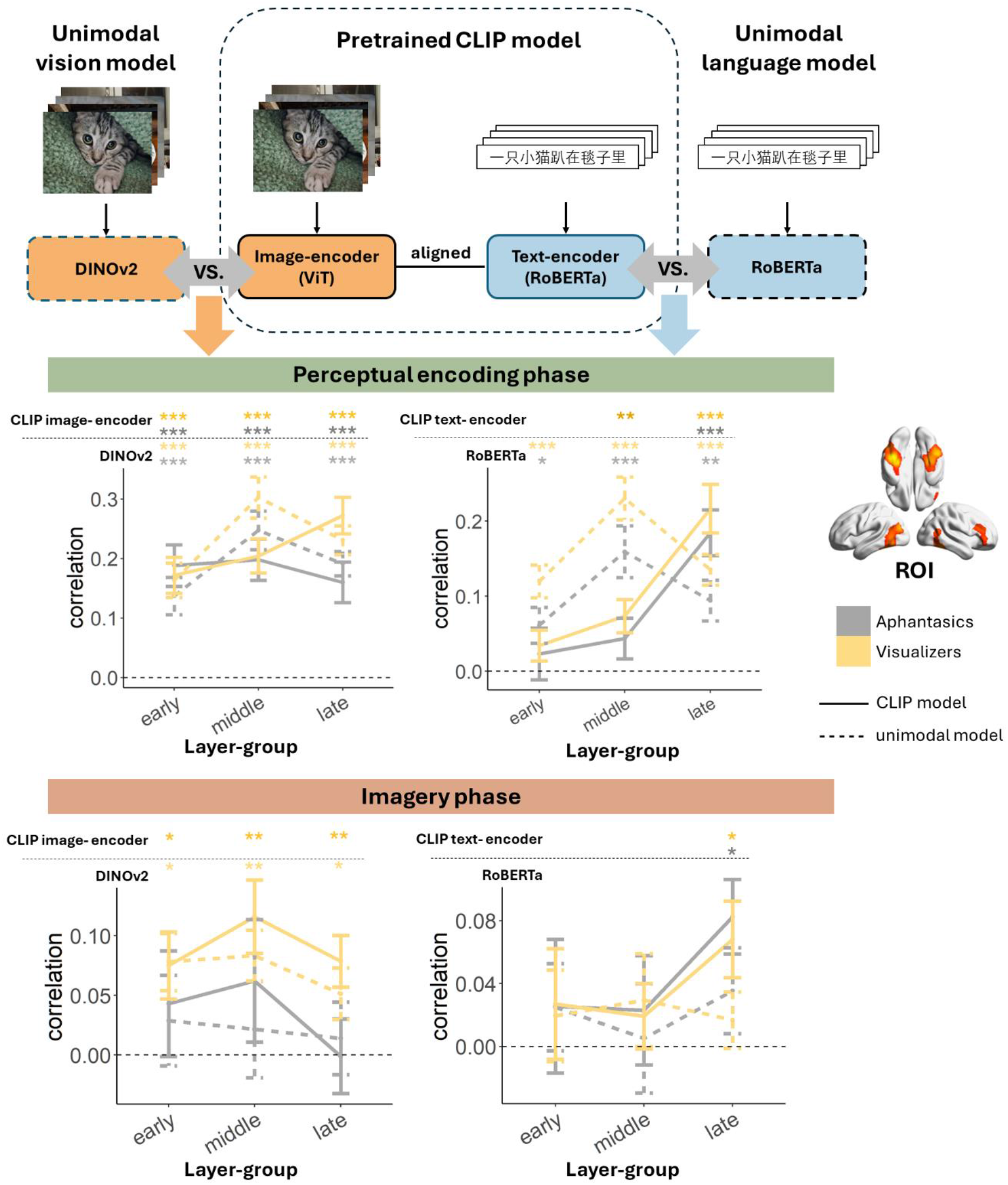
Comparisons between multimodal and unimodal DNNs. Left column: RSA with vision models (DINOv2 and CLIP image-encoder). **Right column:** RSA with language models (RoBERTa and CLIP text-encoder). The ROI was defined from the searchlight SVM cross-phase-decoding map (Fig. 2b). Layers of the vision and language models were grouped into early, middle, and late processing stages (vision: layers 1-8, 9- 16, and 17-24; language: layers 1-4, 5-8, and 9-12). RSA values were averaged within each layer-group, and statistical analyses were performed at the level of these groups. Asterisks indicate significant one-sample t-tests. Significance markers: *p < .05; **p < .01; ***p < .001.

**Fig. 6.**
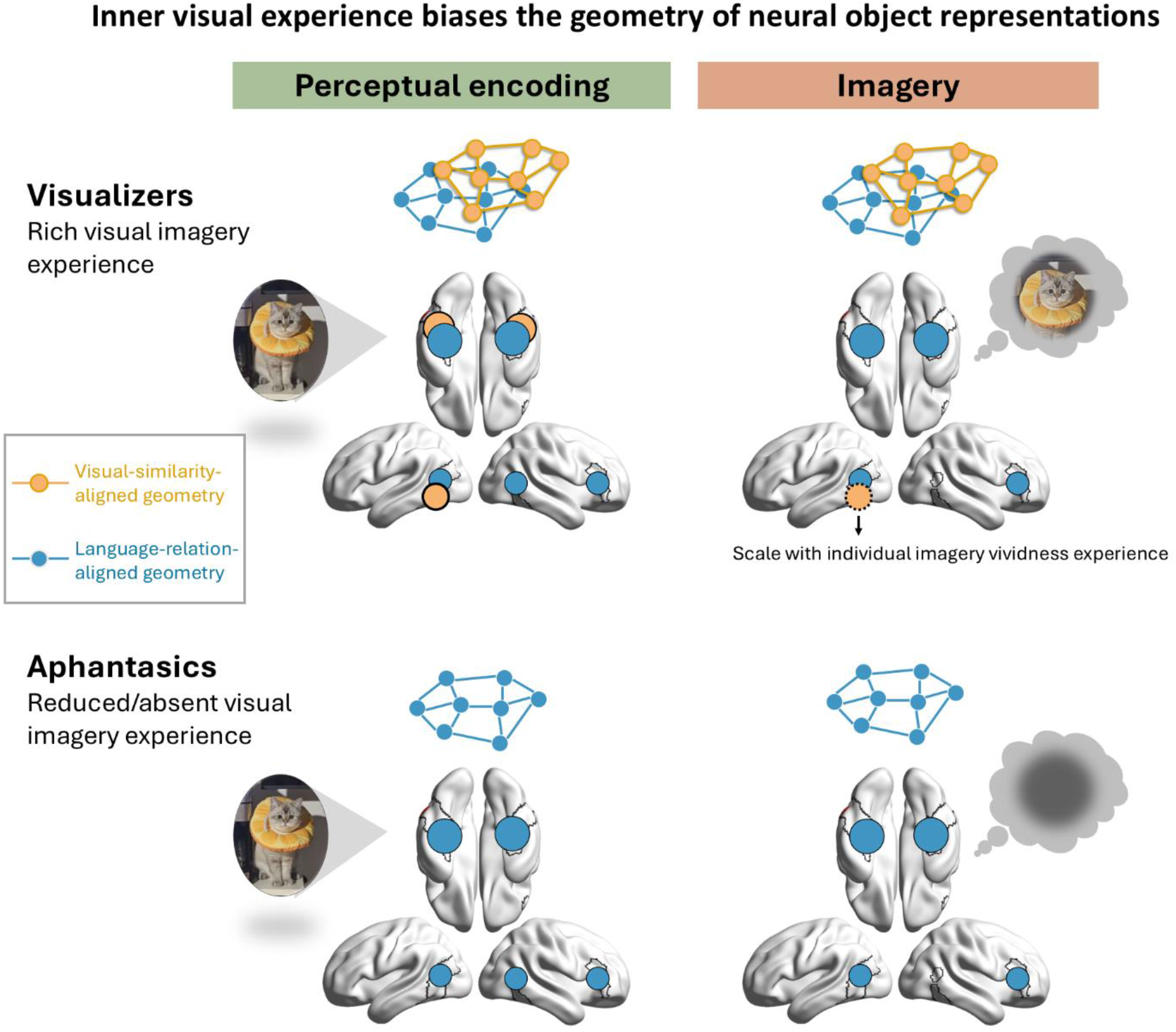
**Summary of how inner visual experience biases the geometry of neural object representations.**Within regions exhibiting perception-imagery cross-decoding (outlined in black), people with and without inner visual experience exhibit distinct representational geometries during both perceptual encoding and imagery. **During perceptual encoding**, **visualizers** with typical inner visual experience encode object information through two types of representational geometries: one uniquely aligned with visual similarity (orange) and the other uniquely aligned with language relations (blue). Visual-similarity-unique representations are distributed across the bilateral fusiform gyrus (FG) and left lateral occipitotemporal cortex (LOTC). Language-relation-unique representations are distributed across the bilateral FG, posterior middle temporal gyrus (pMTG), and right inferior frontal gyrus (IFG). **Individuals with aphantasia**, who lack inner visual experience, encode object information through the representational geometry only uniquely aligned with language-relation. **During imagery**, visual-similarity-unique representations in the left LOTC scale with imagery vividness across all participants. Both visualizers and aphantasics maintain object information through the representational geometry uniquely aligned with language-relation in the bilateral FG, left pMTG, and right IFG.

Specifically, regions exhibiting perception-imagery cross-decoding encompassed multiple representational geometries. Unique visual-similarity-aligned geometry was observed in visualizers across the bilateral fusiform gyrus (FG) and left lateral occipitotemporal cortex (LOTC), with its expression in the left LOTC selectively reduced in individuals with aphantasia during perceptual encoding. The expression of this geometry in the left LOTC during imagery positively tracked subjective imagery vividness across individuals. In contrast, unique language-relation-aligned geometry was preserved across perceptual encoding and imagery in both visualizers and individuals with aphantasia, and was localized to the bilateral FG, left posterior middle temporal gyrus (pMTG), and right inferior frontal gyrus (IFG). Thus, individual differences in visual imagery experience are associated not with the presence or absence of shared neural representations per se, but with selective variation in specific representational geometries.

## Neural geometry uniquely aligned with visual-similarity is associated with imagery experience during both perceptual encoding and imagery phases

The strongest evidence for the representation alteration associated with aphantasia came from the perceptual encoding phase. The neural representations uniquely aligned with image-encoder, i.e., visual-aligned representations, were attenuated in aphantasics relative to visualizers, with the effect localized primarily to the left LOTC. Notably, this difference emerged while both groups were viewing identical visual stimuli, before the imagery phase had even begun. This finding is somewhat counterintuitive. Because the subsequent imagery phase explicitly required participants to generate mental images of the cued object, one might expect imagery-related differences to be most pronounced during imagery itself. Instead, the largest group difference was observed during perception encoding.

These findings suggest that imagery phenotype is associated not only with internally generated representations, but also with the organization of perceptual representations.

This observation has important implications for the imagery literature. Perceptual representations are commonly treated as a relatively stable, stimulus-driven reference against which the format (and content) of imagery representations is evaluated and inferred (e.g., Chang et al., 2025; J. Liu et al., 2025). Our results challenge this assumption. Individuals who differ in subjective imagery experience also differ in the extent to which perceptual representations preserve visual-aligned relational structure. Thus, variation in imagery experience may reflect broader differences in representational organization rather than differences restricted to imagery-specific mechanisms. Importantly, these differences emerged despite comparable behavioral performance, indicating that successful object processing could be achieved by multiple underlying neural representational formats.

The imagery phase results converged with this interpretation. Although categorical group differences were weaker during imagery, the degree to which neural representations uniquely respected image-encoder geometry covaried with subjective imagery vividness. Individuals with more vivid imagery showed stronger alignment with image-based representational structure. Together with the perceptual-encoding findings, these results suggest that imagery experience is specifically associated with the preservation or expression of visual-similarity-aligned object relationships. Thus, imagery experience appears to shape how object knowledge is organized, rather than just an epiphenomenal byproduct of otherwise intact object representations.

## Neural geometry uniquely aligned with language-relation is robust to imagery experience across phases

In contrast to the imagery-related variation observed for visual-aligned geometry, language-relation- aligned geometry was remarkably stable. Significant text-encoder-unique effects were observed during both perceptual encoding and imagery, in both visualizers and aphantasics, and were localized primarily to the bilateral FG, pMTG, and right IFG. Importantly, this does not imply that VOTC representations become linguistic. Rather, it suggests that language-derived relational structure provides a more complete description of the representational geometry that emerges after visual input has been transformed through the ventral visual pathway. Such an interpretation is consistent with converging evidence that language experience contributes to the organization of conceptual representations in visual cortex (Chen et al., 2025; B. Liu et al., 2025), while leaving open the possibility that other sensorimotor experiences also contribute to these higher-order representational structures.

The anatomical distribution of the unique language-relation-aligned effects is broadly consistent with evidence that VOTC regions, especially the FG, support modality-independent object knowledge (Bi et al., 2016; Fairhall & Caramazza, 2013; He et al., 2013; Mahon et al., 2009; Mattioni et al., 2020; van den Hurk et al., 2017; X. Wang et al., 2015). The present findings extend this literature by characterizing the geometry of these representations and demonstrating their preservation across cognitive processes (with or without object visual presence) and populations (with or without mental imagery). This pattern complements findings in congenitally blind individuals in an intriguing way. Congenital blind individuals lack both external-driven visual experience and internally generated visual imagery. They show preserved neural representations in the anterior temporal lobe about vision-only-knowledge such as color, presumably a kind of language-relation-aligned representation (Bottini et al., 2020; Striem-Amit et al., 2018; X. Wang et al., 2020). However, their VOTC neural representation for such knowledge, was absent, without evidence of language-aligned representation in VOTC (X. Wang et al., 2020), unlike aphantasics. Thus, external-driven visual perception experience seems to be critical for the formation of language-relation-aligned representational geometry in visual cortex.

The pMTG is a region identified to be part of the language network (Binder et al., 2009; Binder & Desai, 2011; Fairhall & Caramazza, 2013; Jackson, 2021; Ralph et al., 2017; Tune & Asaridou, 2016), and has been implicated in multimodal semantic processing and semantic control (Y. Xu et al., 2016). Its partial overlap with LOTC, which exhibited imagery-related visual-similarity-unique representations, suggests that it may flexibly integrate modality-specific and modality-independent information depending on task demands. The right IFG, often associated with working memory and control processes, may reflect the engagement of internally maintained semantic representations that are not strictly tied to the modality of external input (Koelsch et al., 2009; Kumar et al., 2016; Schneiders et al., 2012; Shivde & Thompson-Schill, 2004; Uluç et al., 2018).

Considered together, the differential relationships between imagery experience and visual- vs. language- relation- aligned representational geometries have broader implications for how perception-imagery overlap is interpreted. The regions analyzed here were selected precisely because they showed successful cross-perception-imagery decoding for both subject groups, a standard operational definition of shared representations in the imagery literature (Dijkstra et al., 2017; Harrison & Tong, 2009; Lee et al., 2012; Reddy et al., 2010). However, decoding success alone is largely agnostic about representational content. Indeed, within regions showing reliable cross-phase-decoding, neural representations differed across groups (and across phases) in the extent to which they preserved unique visual-aligned structure. Thus, successful correspondence between perception and imagery can arise from representational spaces that are only partially shared. More generally, the notion of a shared perception-imagery code may require refinement. Rather than reflecting a single common representational format, perception and imagery are supported by heterogeneous representational geometries that reflect multiple organizational principles. Different representational geometries may vary in their expression across individuals, cognitive states, and brain regions while still supporting successful object processing. Characterizing representational geometry therefore provides information that cannot be recovered from activation overlap or decoding analyses alone.

## Implications for aphantasia and the functional imagery node

Previous studies reported that individuals with aphantasia showed reduced neural similarity between perception and imagery within a specific fusiform region, which was coined the “fusiform imagery node” (FIN; J. Liu et al., 2025; Spagna et al., 2024), as well as in the early visual cortex (Chang et al., 2025).

Although our whole-brain cross-perception-imagery-decoding analysis did not reveal statistically significant group differences after correction for multiple comparisons, it showed a trend toward reduced cross-decoding in regions adjacent to the FIN (Supplementary Figs. S2-3). In contrast, we found no reliable cross-decoding in EVC in either group, in line with broader literature that shared representations between object perception and imagery are primarily supported by higher-level visual cortices (see Pearson, 2019; Pearson & Kosslyn, 2015; Spagna et al., 2024 for comprehensive discussions of the role of EVC in visual imagery).

Importantly, our representational content analyses uncovered group differences extended beyond the FIN to more lateral and dorsal regions, spanning to the pMTG. This is naturally explained by the discussion earlier that cross-perception-imagery-decoding may underestimate the changes in aphantasia: the preservation of language-relation-aligned representations may be sufficient to support successful cross-phase decoding even when visual-aligned representations are diminished. Together with the widespread absence of representations uniquely aligned with visual-similarity across VOTC in individuals with aphantasia, these findings suggest that visual imagery may associate less with a discrete functional "imagery node" than on distributed visual-aligned representations spanning the higher visual cortex.

This identified left LOTC has been found supporting a broad range of conceptual knowledge, including representations of tools (Chao et al., 1999; Martin et al., 1996), actions (Karakose-Akbiyik et al., 2023; Moritz et al., 2015), and object shapes (Y. Xu et al., 2023). Notably, many of these representations are preserved in congenitally blind individuals despite the absence of visual experience (Bedny et al., 2012; Dormal et al., 2018; Y. Xu et al., 2023; see Wurm & Caramazza, 2022 for a review), highlighting the multimodal nature of this region. Unlike congenitally blind individuals, who may manage to form non-visual sensory imagery (e.g., haptic, taste, smell, sound), most individuals with aphantasia report reduced imagery across multiple sensory modalities (Dance et al., 2021). From this perspective, the altered representations observed in the left LOTC may reflect the absence of multimodal mental imagery rather than visual imagery alone. Nevertheless, because non-visual imagery abilities were not systematically assessed in the present study, this possibility remains speculative and should be examined directly in future work.

## Multimodal alignment is critical in the absence of visual input

Our comparison between multimodal and unimodal models also provides insight into the representational principles underlying VOTC activity. Previous work has shown that, during visual object perception, visual representations from language-supervised vision-language models (i.e., the CLIP image encoder) uniquely account for VOTC activity when regressing out representations from unsupervised vision models such as MoCo, suggesting that language supervision reshapes visual representations in higher visual cortex (Chen et al., 2025). The present findings extend this idea one step further. Within the same multimodal framework, neural representational geometry in VOTC was even more robustly explained by the CLIP text-encoder than by the CLIP image-encoder when either external visual input was absent (imagery) or internal visual experience was unavailable (aphantasia). This finding suggests that VOTC contains not only visual representations shaped by language supervision, but also higher-order relational representations directly encoded in language itself.

The direct comparison between multimodal and unimodal models further revealed that the explanatory advantage of multimodal alignment depended on cognitive state. During perceptual encoding, unimodal vision and language models accounted for much of the neural variance and even outperformed their corresponding CLIP encoders in some processing stages, suggesting that sensory-driven processing remains strongly constrained by modality-specific statistics. During imagery, however, both CLIP encoders showed a relative advantage over their unimodal counterparts. Although these differences were modest, they suggest that when representations must be internally maintained in the absence of visual input, representational spaces jointly shaped by visual and linguistic experience become increasingly informative. More generally, together with previous comparisons across perception tasks, these findings suggest that the correspondence between computational models and brain representations is itself dynamic, varying not only with model architecture and training objective, but also with cognitive state and task demands. Such flexible representational principle may provide a neural substrate that supports access to object knowledge across diverse perceptual and cognitive situations.

To conclude, perceptual and imagery experiences are coupled in ways different from previously thought. Visual-similarity-aligned neural geometry in perceptual encoding was already associated with imagery experience. Language-aligned relational structure remained stable across imagery phenotypes and processing phases. These findings suggest that subjective imagery experience is associated not merely with imagery-specific processes, but with how the brain encodes objects for perception and knowledge more broadly.

## Methods Participants

Seventeen healthy individuals with congenital aphantasia (mean age ± SD: 23 ± 3 years; 14 females) and twenty-two control participants with typical visual imagery (22 ± 2 years; 17 females) took part in the study. Aphantasia was identified using the Vividness of Visual Imagery Questionnaire (VVIQ, Marks, 1973), with scores below 32. Follow-up interviews ensured participants fully understood the instructions and confirmed the absence or near absence of visual imagery experience. One participant with a VVIQ score of 35 was also included in the aphantasic group, as their subjective report closely aligned with the characteristics of aphantasia. All participants had normal or corrected-to-normal vision and hearing, and no history of psychiatric or neurological disorders. Informed consent was obtained from all participants, who received monetary compensation. The study was approved by the Institutional Review Board of the State Key Laboratory of Cognitive Neuroscience and Learning at Beijing Normal University, and adhered to the Declaration of Helsinki.

## Stimuli

Three categories of images were used (Supplementary Fig. 1): two meaningful domains—animals (e.g., cat, tortoise, elephant, crab, panda, bee) and large objects (e.g., tea table, dressing table, bathtub, fence, sailboat, children’s slide)—and one meaningless shape class. The meaningless shape class was not of interest in the current study and was therefore not included in the analysis.

All images were converted to grayscale and normalized for luminance and contrast using the SHINE toolbox (Willenbockel et al., 2010). Familiarity and pixel-based inter-image dissimilarity between animal and large object images were matched (two-sample t-test ps > .08). The test images (used in the test phase; see Procedure) were visually similar but distinct exemplars.

## Procedure

We employed a retro-cue paradigm (Fig. 1c; Dijkstra et al., 2017; Harrison & Tong, 2009). In each trial, two images were shown sequentially for 1 s each (perception phase), separated by a jittered interstimulus interval of 2 - 4 s. A numeric cue (‘1’ or ‘2’) then indicated which image participants should visualize during a subsequent 4 s blank interval (imagery phase). Finally, a test image was presented (test phase), and participants judged whether it was exactly the same image, not just the same object, as the cued one. Responses were made with the index fingers (left = "yes", right = "no"). Each trial lasted 17 s.

To optimize the experimental design, trial sequences were counterbalanced: the two sequentially presented stimuli always came from different categories, each had an equal probability of appearing first or second, and each was equally likely to be cued. Image pairings were unique across every four runs. The numbers of “yes” and “no” trials were also balanced. Each run contained 18 trials, such that within each run every item was imagined once and perceived twice.

Thirty-seven participants completed all 8 runs; two (one from each group) completed only 4 runs due to personal reasons.

## Image acquisition

MRI data were collected on a Siemens Prisma 3T scanner using a 64-channel phased-array head coil at Beijing Normal University Neuroimaging Center. Functional images were acquired with a multi-band echo-planar imaging sequence: 64 axial slices, 2 mm thickness, phase encoding direction from posterior to anterior; 0.2 mm gap; multi-band factor = 2; TR = 2000 ms; TE = 30 ms; flip angle = 90°; matrix size = 104 × 104; FoV = 208*208; voxel size = 2 × 2 × 2 mm^3^. Each task run lasted 5 minutes and 34 seconds (167 volumes). High-resolution T1-weighted anatomical images were collected with a 3D MPRAGE sequence: 192 sagittal slices; 1 mm thickness; TR = 2530 ms; TE = 2.98 ms; TI = 1100 ms; flip angle = 7°; FoV = 224 × 256 mm^2^; voxel size = 0.5 × 0.5 × 1 mm^3^, interpolated; matrix size = 224 × 256.

## Data preprocessing

Functional data preprocessing was conducted in SPM12 (<u>SPM12 Software - Statistical Parametric Mapping</u>) using MATLAB R2020a (v.2020a, MathWorks). The first 6 volumes, acquired during the initial fixation period, were discarded. The remaining data underwent slice-timing correction, motion correction, and high-pass filtering (cutoff: 0.008 Hz). Data were normalized to Montreal Neurological Institute (MNI) space using unified segmentation and resampled to 2 mm isotropic voxels

## Data analysis

### Behavioral Performance

Accuracy was calculated separately for each stimulus class and each participant, followed by a repeated-measures ANOVA with group and stimulus class as factors. For reaction times (RT), we included only correct trials and excluded outliers beyond ±3 standard deviations. RTs were then averaged across trials for each participant and class, followed by another repeated-measures ANOVA with the same factors.

### GLM Construction

We modeled each subject’s data using a general linear model (GLM) for each run in SPM12. Predictors of interest consisted of separate regressors for each item identity during the perception, imagery, and test phases, with six motion parameters included as nuisance regressors. All predictors were convolved with the canonical hemodynamic response function. Since the test phase was only used to ensure task engagement, corresponding data were excluded from further analyses. For multivariate decoding analyses, beta images were spatially smoothed using a 2-mm FWHM Gaussian kernel (Hendriks et al., 2017; Op de Beeck, 2010).

### Searchlight SVM Decoding Analysis

Searchlight analysis was restricted to a whole-brain gray matter mask, defined as voxels with probability > 1/3 in the SPM12 gray matter template and located within cerebral regions (1 - 90) of the Automated Anatomical Labeling (AAL) atlas, yielding 122,694 voxels (981,552 mm³). For each voxel, we defined an 8-mm radius sphere (257 voxels) and extracted beta estimates for each item and run, z-score normalized within the sphere. We employed repeated split-half cross-validation (Valente et al., 2021), exhaustively iterating through all possible data splits, yielding 70 unique split-half partitions for participants with 8 runs and 6 combinations for those with 4 runs. In each split, perception data from one half were used to train an SVM classifier, which was then tested on imagery data from the other half, and vice versa. The mean accuracy across both directions and all splits was assigned to the sphere’s center voxel. Group-level statistical inference was performed using permutation-based Statistical nonParametric Mapping (SnPM) (<u>NISOx: SnPM</u>), with 5000 permutations and no variance smoothing. All searchlight decoding results were thresholded at voxelwise p < .001, cluster-level FWE-corrected p < .05, unless stated otherwise. Domain-level decoding classified animals vs. large objects (chance-level: 1/2); item-level decoding classified among individual object identities (chance-level: 1/12).

### Features extraction from DNN models

***CLIP*** is a neural network trained on a variety of image-text pairs, being adapted by the alignment between images and their corresponding text descriptions (Radford et al., 2021). We utilized a Chinese version of CLIP (Yang et al., 2023), pretrained on around 200 million publicly available image-text pairs data in Chinese, using the ViT-L/14 architecture for the image-encoder and RoBERTa-wwm-Base for the text-encoder. For comparison, we included unimodal transformer models with closely matched backbones. The vision model was a Vision Transformer (***ViT***) pretrained on LVD-142M using the self-supervised DINOv2 method (Oquab et al., 2024). The language model was a Chinese ***RoBERTa*** model pretrained on Chinese Wikipedia and extended corpora (Cui et al., 2021). Both unimodal models share closely matched transformer backbones with their corresponding CLIP encoders.

For the construction of representational dissimilarity matrix (RDM), we extracted the [CLS] token representations rather than applying mean pooling. RSA compares representational geometry across whole objects, requiring a single embedding to represent each stimulus. We therefore used the model’s canonical sequence-level representation, namely the [CLS] token, which is explicitly optimized during training to summarize the entire input into a global object representation. In contrast, mean pooling constitutes an additional aggregation strategy applied after the forward pass rather than one directly optimized by the model’s training objective, and may alter the representational geometry by averaging contextualized patch embeddings. To obtain layer-wise representations, we registered forward hooks on the output of each transformer block in the visual models (i.e., the CLIP-image encoder and DINOv2) and on each layer of the BERT-based text models (i.e., the CLIP text-encoder and RoBERTa). During a forward pass, we collected the [CLS] token embeddings from each layer for both image and text inputs.

Experimental stimuli (preprocessed images) were passed through the visual encoders, whereas object names (e.g., “cat,” “elephant,” “dressing table” in Chinese) were tokenized and fed into the text encoders. At each layer, [CLS] embeddings were detached, stored, and stacked across stimuli. Cosine similarity matrices were computed from L2-normalized features and converted into RDMs by subtracting similarity values from one. Layer-wise correlations between models of the same modality are shown in Supplementary Fig. 7.

For CLIP model, although the contrastive objective influences the entire encoder via backpropagation, its impact is expected to intensify toward the final layers that are directly optimized for cross-modal alignment. The penultimate layer thus offers a principled balance between preserving modality-biased structure and maintaining task-relevant high-level representations. Accordingly, the penultimate layer was used in the primary analyses.

When comparing multimodal and unimodal models, we additionally considered potential differences in the hierarchical emergence of high-level representations across architectures. Because these models are optimized with distinct training objectives (Kornblith et al., 2021; Yamins & DiCarlo, 2016; Yosinski et al., 2014), abstract and object-level information may arise at different depths within the network.

Therefore, rather than restricting comparisons to a single layer, we examined representational similarity across multiple processing stages. To ensure interpretable and statistically stable comparisons, and to avoid over-fragmentation across individual layers, we grouped layers into early, middle, and late stages. For the vision models, the early, middle, and late stages comprised layers 1-8, 9-16, and 17-24, respectively. For the language models, the corresponding stages comprised layers 1-4, 5-8, and 9-12.

This approach captures broad hierarchical patterns without introducing unnecessary complexity in the presentation and interpretation of results. Layer-wise RSA values were averaged within each layer-group, and statistical tests were performed at the layer-group level.

### Linear mixed-effects model analysis of RSA with CLIP encoders

Neural RDMs were constructed using t-value vectors extracted from the ROI defined based on the searchlight SVM result (see Results and Fig. 2b). For each subject, the off-diagonal entries of the neural RDM were vectorized and each pairwise dissimilarity constituted one observation. Corresponding entries from the CLIP image-encoder and text-encoder RDMs were used as predictors. Linear mixed-effects models were fitted separately for the perceptual encoding and imagery phases using restricted maximum likelihood estimation. Subject was included as a random effect to account for non-independence of observations within individuals. The fixed effects included CLIP image-encoder RDM (derived from the penultimate layer), CLIP text-encoder RDM (derived from the penultimate layer), group, and their interactions. Random slopes were included where supported by the data to account for individual variability in representational sensitivity. The models were specified as follows:

Perceptual encoding phase:

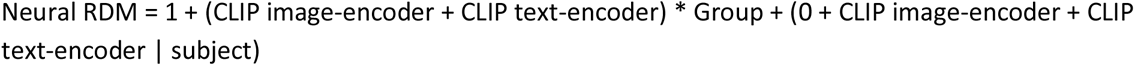

Imagery phase:

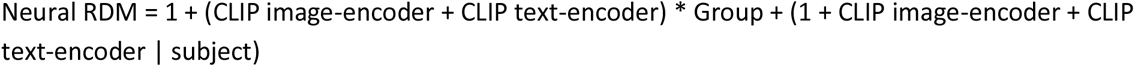

The final random-effects structure was determined based on model convergence and singularity diagnostics.

### Searchlight RSA with CLIP encoders

Searchlight RSA was conducted within the cross-phase-decoding mask (see Results and Fig. 2b). Neural RDMs were computed for each subject by extracting T-value vectors from searchlight spheres (as in the decoding analysis), and measuring dissimilarities as 1 − Pearson correlation between stimulus pairs. We then computed Spearman partial correlations between neural RDMs and one encoder RDM, controlling for the corresponding RDM from the other encoder.

Coefficients were assigned to the center voxel of each sphere. Group-level analyses were also performed using permutation-based Statistical nonParametric Mapping (SnPM) (<u>NISOx: SnPM</u>), with 5000 permutations and no variance smoothing. Results were thresholded at voxelwise p < .001, cluster-level FWE-corrected p < .05, unless stated otherwise.

### Statistical tests statement

*ROI-based* statistical analyses were conducted in R (R Core Team, 2019) using the rstatix package (Kassambara, 2021). All correlation coefficients used for inferential statistics were Fisher z-transformed prior to analysis. Linear mixed-effects models were fitted using the lme4 package (Bates et al., 2015), with p-values estimated via the lmerTest package (Kuznetsova et al., 2017). Two-sample comparisons were conducted using Welch’s t-test, which adjusts for unequal variances and sample sizes (Delacre et al., 2017). Multiple comparisons were controlled using false discovery rate (FDR) correction where applicable.

## Acknowledgement

We thank Rui Feng, Jiahui Lu, Haojie Wen, and Ziyi Xiong for their assistance with data collection; Ze Fu for his advice on the analysis; Haoyang Chen for his assistance with the utility of the DNN models; Ze Fu, Haojie Wen, and Huichao Yang, for their advice on the figures; Yiyang Cai, Ziyi Xiong, and Xiaosha Wang for their advice on the manuscript.

## Data availability

CLIP models were acquired from: https://github.com/OFA-Sys/Chinese-CLIP.

DINOv2 were acquired from: https://huggingface.co/timm/vit_large_patch14_reg4_dinov2.lvd142m.

RoBERTa were acquired from: https://github.com/ymcui/Chinese-BERT-wwm.

The data supporting the findings of this study, including behavioral data, fMRI statistical maps, and representational similarity matrices derived from ROI analyses, will be deposited in a public repository and made available upon publication. All data have been fully anonymized in accordance with institutional and ethical guidelines.

## Declaration of generative AI use

During the preparation of this work the authors used ChatGPT (OpenAI) to assist with language and grammar editing. After using this tool, the authors reviewed and edited the content as needed and take full responsibility for the final manuscript.

