## Supplementary for "Inner visual experience biases object neural representational geometry during perceptual encoding"

### Supplementary Materials

**Supplementary Table 1**

**Supplementary Figures 1-7**

**Table 1.** Results of three-way repeated-measures ANOVA

**Perceptual encoding phase, vision models**

| Effect | Df | F | p | $\eta2_p$ |
|---|---|---|---|---|
| subject-group | 1,37 | 1.03 | 0.32 | 0.03 |
| layer-group | 1,37 | 16.58 | **$1.91\times10^{-5}$** | 0.31 |
| model | 2,74 | 4.41 | **0.04** | 0.11 |
| subject-group × layer-group | 2,74 | 4.32 | **0.03** | 0.11 |
| subject-group × model | 1,37 | 0.47 | 0.50 | 0.01 |
| layer-group × model | 2,74 | 9.97 | **0.001** | 0.21 |
| Subject-group × layer-group × model | 2,74 | 4.00 | **0.04** | 0.10 |

**Perceptual encoding phase, language models**

| Effect | Df | F | p | $\eta2_p$ |
|---|---|---|---|---|
| subject-group | 1,37 | 2.00 | 0.17 | 0.05 |
| layer-group | 1,37 | 26.34 | **$1.20 \times10^{-7}$** | 0.42 |
| model | 2,74 | 7.87 | **0.008** | 0.18 |
| subject-group × layer-group | 2,74 | 0.19 | 0.77 | 0.005 |
| subject-group × model | 1,37 | 1.36 | 0.25 | 0.04 |
| layer-group × model | 2,74 | 39.20 | **$2.46 \times10^{-12}$** | 0.51 |
| Subject-group × layer-group × model | 2,74 | 0.47 | 0.63 | 0.01 |

**Imagery phase, vision models**

| Effect | Df | F | p | $\eta2_p$ |
|---|---|---|---|---|
| subject-group | 1,37 | 1.70 | 0.20 | 0.04 |
| layer-group | 1,37 | 2.54 | 0.10 | 0.06 |
| model | 2,74 | 4.66 | **0.04** | 0.11 |
| subject-group × layer-group | 2,74 | 0.24 | 0.75 | 0.003 |
| subject-group × model | 1,37 | 0.09 | 0.76 | 0.003 |
| layer-group × model | 2,74 | 1.12 | 0.32 | 0.03 |
| Subject-group × layer-group × model | 2,74 | 0.89 | 0.38 | 0.02 |

**Imagery phase, language models**

| Effect | Df | F | p | $\eta2_p$ |
|---|---|---|---|---|
| subject-group | 1,37 | 0.02 | 0.89 | $4.96\times10^{-4}$ |
| layer-group | 1,37 | 1.96 | 0.16 | 0.05 |
| model | 2,74 | 1.39 | 0.25 | 0.04 |
| subject-group × layer-group | 2,74 | 0.30 | 0.68 | 0.008 |
| subject-group × model | 1,37 | 0.05 | 0.83 | 0.001 |
| layer-group × model | 2,74 | 1.36 | 0.26 | 0.035 |
| Subject-group × layer-group × model | 2,74 | 0.20 | 0.82 | 0.005 |

**Fig. S1| Stimuli**

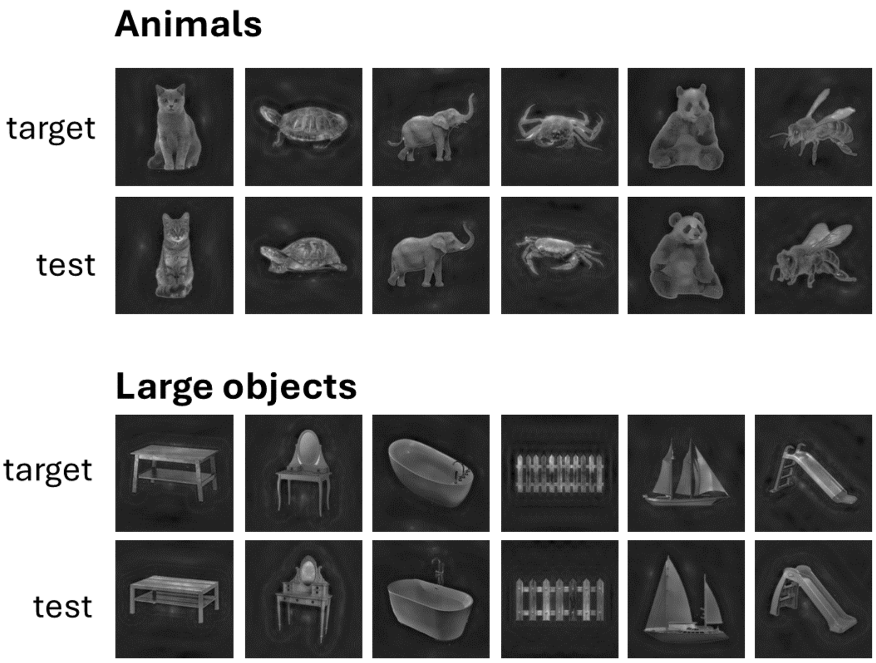

**Fig. S2 | Searchlight SVM cross-phase decoding for object domains (animal vs. large object) in the visualizer group and aphantasic group separately**

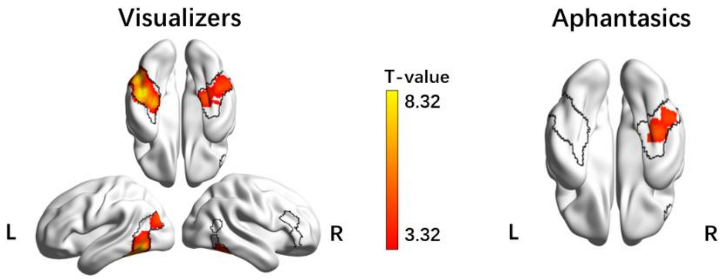


*p < .001, cluster-level FWE-corrected p < .05*

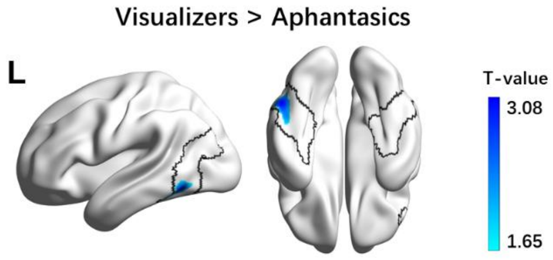


*p < .05, uncorrected*

**Fig. S3 | Searchlight SVM cross-phase decoding for object identities (12-way)**

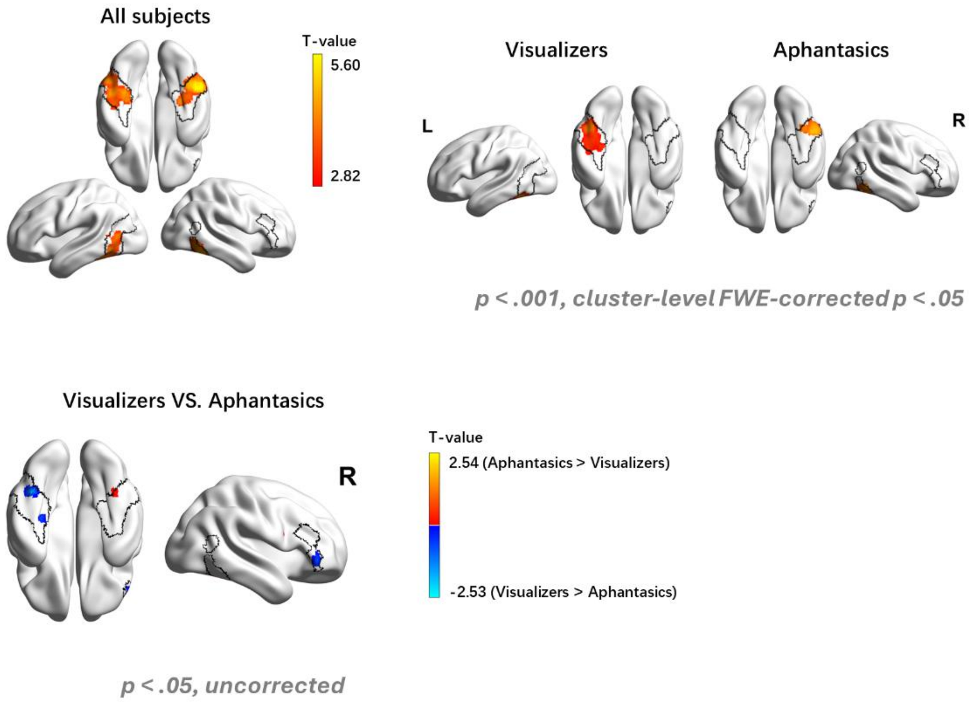

**Fig. S4 | Uncorrected results of searchlight RSA with CLIP encoders (partial-correlation-based)**

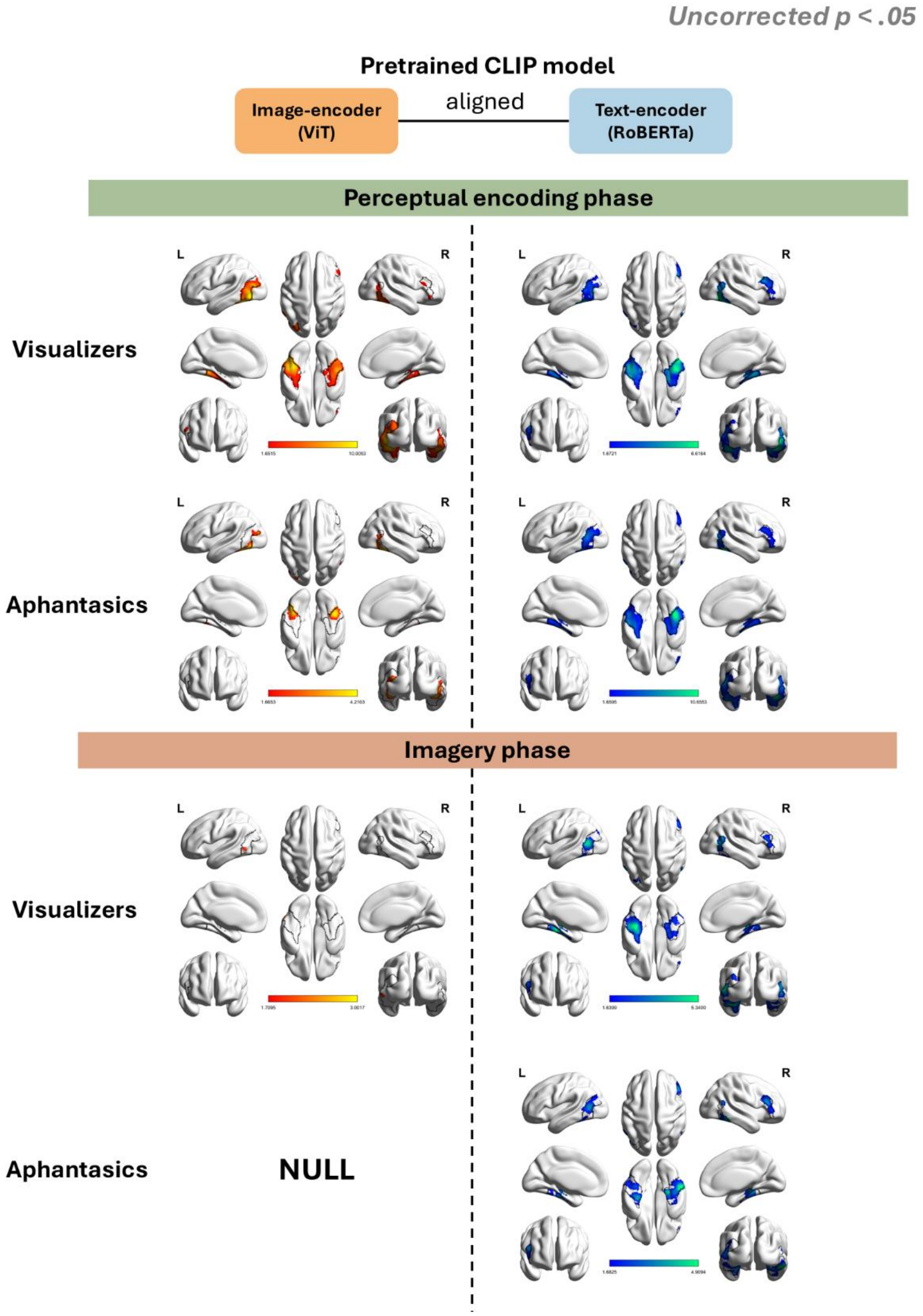

**Fig. S5 | Uncorrected results of group comparison in searchlight RSA with CLIP encoders (partial-correlation-based)**

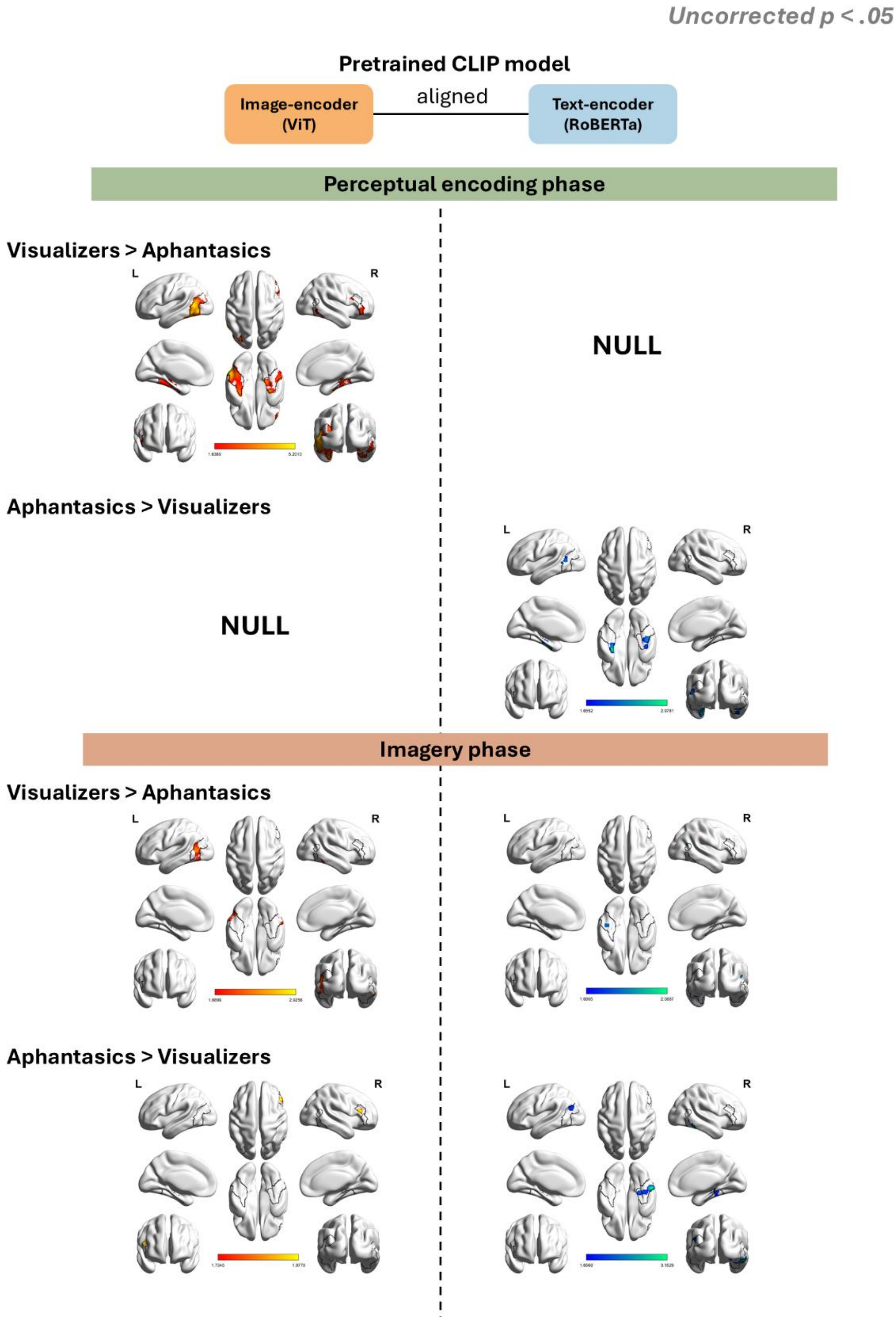

**Fig. S6 | Searchlight RSA with CLIP encoders (zero-order-correlation-based & commonality-based)**

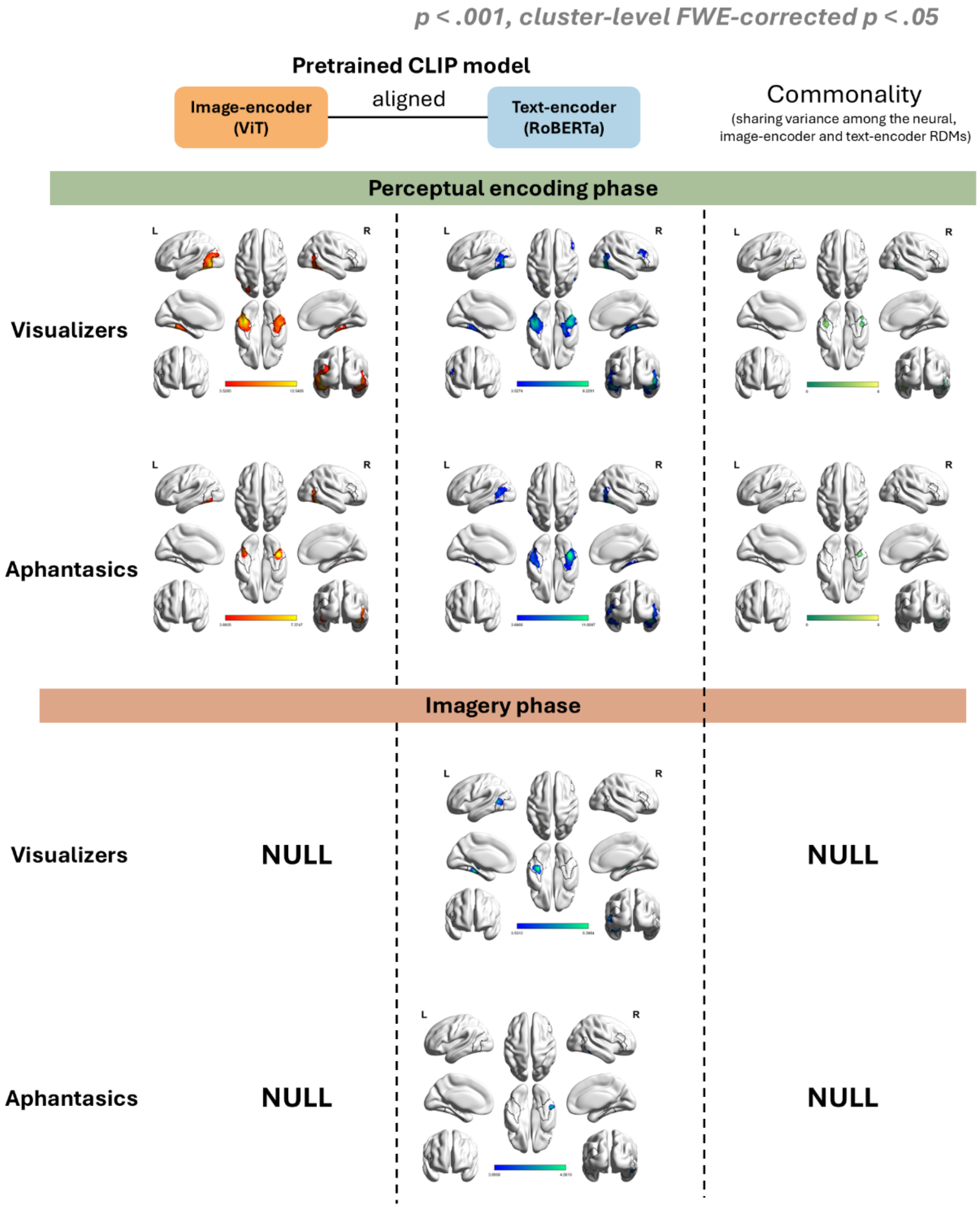

**Fig. S7 | Heatmaps of correlations between layers of DNN models**

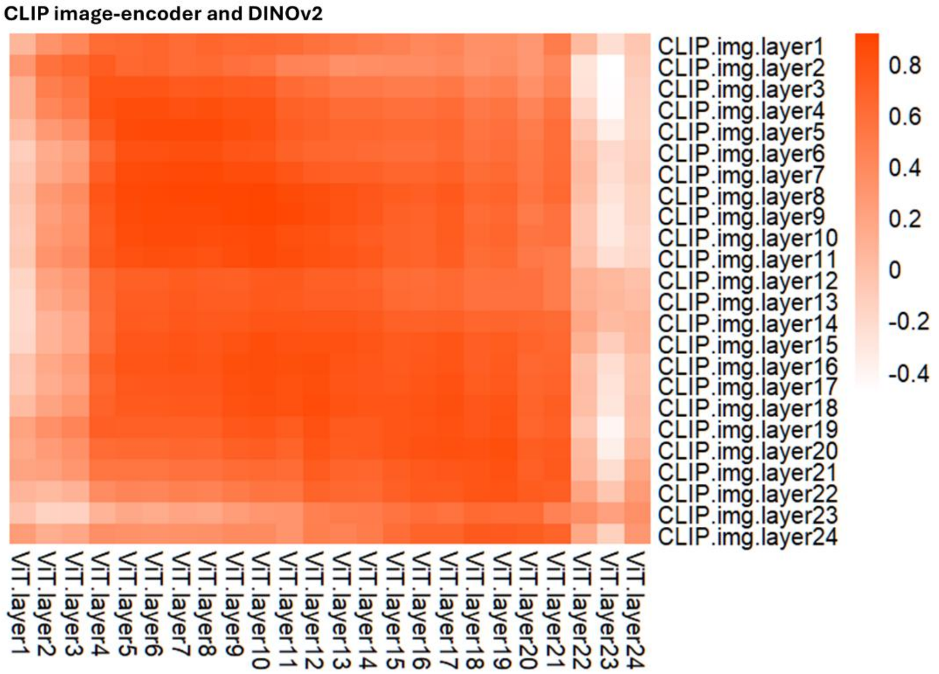


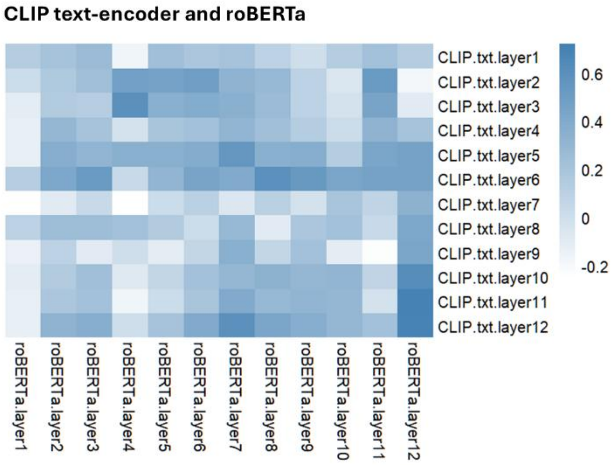
